# On the effects of climate change on Galápagos green turtle *Chelonia mydas agassizii* neonate mortality in A1B

**DOI:** 10.1101/2020.02.27.968628

**Authors:** Max Gotts

## Abstract

Climate change affects every crevice and corner of the ecological world. Already we see seasonal effects on important ecosystems such as the marine habitats of the Galápagos Archipelago. This paper looks at the year 2100 in the IPCC scenario A1B, looking specifically at *Chelonia mydas agassizii* and seeing whether it will be a “winner” or a “loser” under climate change. We discover that while neonate hatchlings may benefit from a compressed laying period that offers them protection by way of numbers, and speed from the increased ocean temperatures (SST), they lose out many other places. Frigatebird and reef fish predation will increase, hatchling size and fitness may decrease, and a surplus of female hatchlings will be produced, or none at all as nest temperatures rise above the thermal mortality limit. *Chelonia mydas agassizii* will need serious conservation help if we wish to keep it on this earth by the year 2100 even if we opt for the A1B strategy.

## 1 Introduction

Climate change will indisputably change the lives of almost every creature on Earth. Whether we choose to act now or in 15 years or never at all may change how significant the impacts of climate change are.

Galápagos is a key subject of interest in this regard because many of the species there are endemic, the currents that supply the archipelago are diverse, and because the Galápagos-Cocos sea mounts form a significant rest stop for migrating sharks and marine megafauna.

The Galápagos Archipelago is located at a latitude of −0.47°, and a longitude of −90.4°. It is one of the most significant rookeries for *Chelonia mydas* (green sea turtles) in the world (Zárate 2013). Specifically, the local subspecies is *C. m. agassizii*, coloquially known as the black turtle or Galápagos green turtle. This paper looks at neonate mortality as a function of emergence, oviposition timing, and fitness as a result of nest and water temperature.

Climate change is a daunting issue for any biologist interested in marine chelonians as they are extraordinarily affected by temperature changes. Sea turtles’ sexes are temperature-dependent (TSD)—the temperature of the nest determines the sex of the individuals inside the nest. This makes a warming globe a distressing prospect.

This paper looks into the IPCC scenario A1B, in which the world exists in a balance of fossil-fuel and renewably driven energy-generation systems. We believe that this is the most likely scenario, unless a drastic and incredible change occurs within the next five to ten years. In this paper, we look into the future to 2100 and attempt to analyse what world black turtles will live in, and how their numbers might adjust.

Already, Wolff (2010) and others have shown how climate change is affecting Galápagos. However, the effects almost a century into the future are more diverse (and harrowing) than the effects we already see today.

## 2 Methods

We ran simulations using the EdGCM software, obtaining data for the Predicted SST simulation and the IPCC’s A1B simulation. A1B is the most probable scenario in our opinion given politician’s resistance to approving bills that counteract climate change.

To compare the data, we used the EdGCM app “Panopoly,” and performed a simple subtraction of the arrays. Namely, we looked at cloud cover, sea surface temperature (SST), and surface air temperature (AT). In every case we subtracted A1B variables from Predicted SST variables, and centered the data on 0 so that we could see which areas would experience a positive change and which areas would be affected by a negative change. For all variables, red correlates with increase, and blue correlates with decrease.

We looked at winter, spring, and annual cases for each variable. Annual cases sometimes showed contradicting trends in variable change than winter and spring cases, both in comparison to Predicted SST, which corroborates Wolff (2010)’s claim that whilst Galápagos (in 2010) did not show evidence of warming, it exhibited seasonal changes as a direct result of climate change.

Winter and spring were looked at because these are the seasons that emergence and oviposition usually occur in. Oviposition is consistently 70 days before emergence, and accurate within a week. Zárate *et al*. (2013) puts emergence between the months of December to July. Oviposition, of course, is normally distributed across these months, so it does not seem unreasonable to look primarily at these months for both emergence and oviposition.

## 3 Results Discussion

### 3.1 Cloud Cover

As discussed in the **Methods** section, there are some figures in which the annual contradicts the seasonal data. Cloud cover is one such example.

Notice how in *Fig. 1* cloud cover decreases over the archipelago. However, for the seasons we are interested in, the opposite is true, as shown in *Figure 2*.

**Fig. 1.**
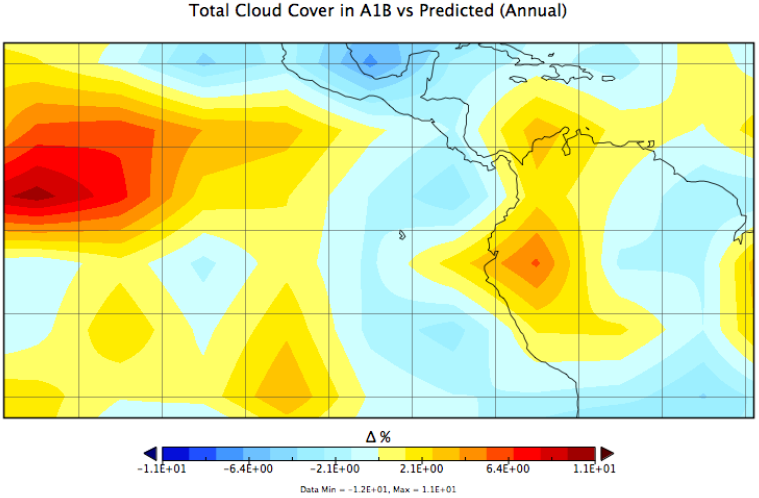
Change in cloud cover (in percentage) between A1B and Predicted SST over Galápagos annually using a equirectangular map projection.

**Fig. 2.**
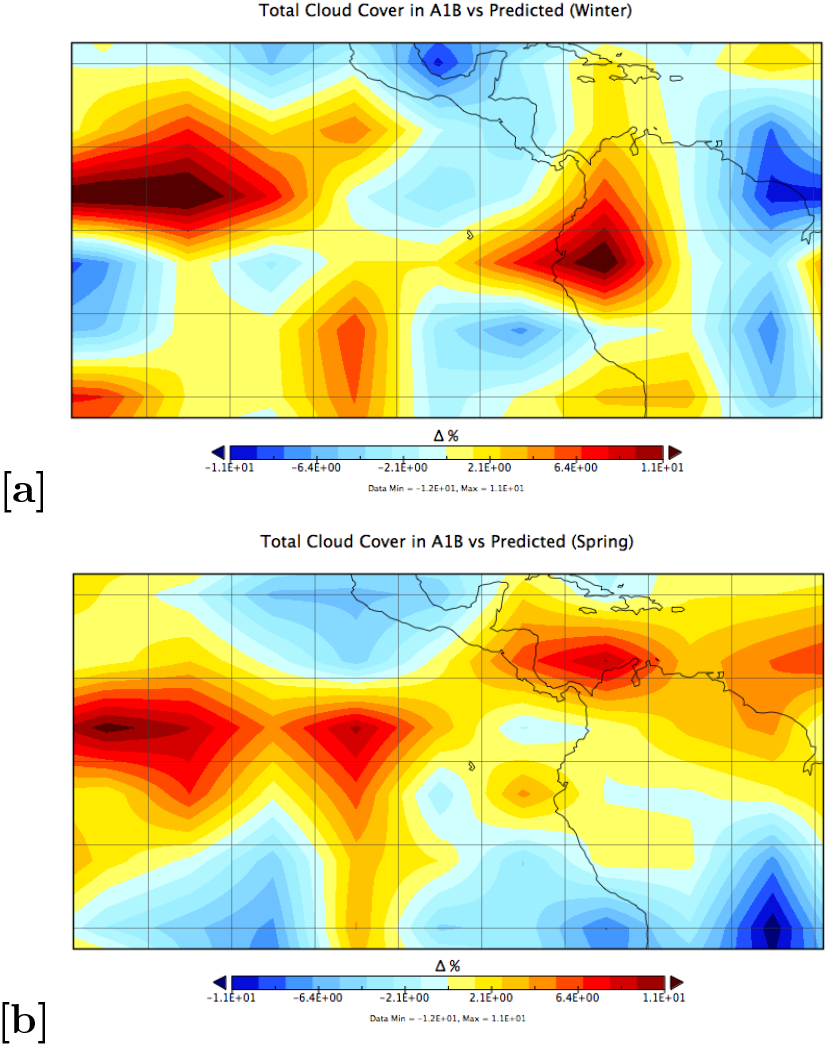
Change in cloud cover (in percentage) between A1B and Predicted SST over Galápagos in **[a]** winter and **[b]** spring using a equirectangular map projection.

In *Fig. 2a, b*, it becomes clear that during the “important” seasons for this analysis cloud cover increases on the order of ≈ 2%.

### 3.2 Surface Air Temperature

Analysing surface air temperature is significantly easier than cloud cover, since the annual accurately represents the winter and spring conditions, as shown below.

The 3° – 4°C difference in *Fig. 3* is a significant change, as we will discuss in the section. However, for now it is enough to note that Fig. 3 matches *Fig. 4a, b*.

**Fig. 3.**
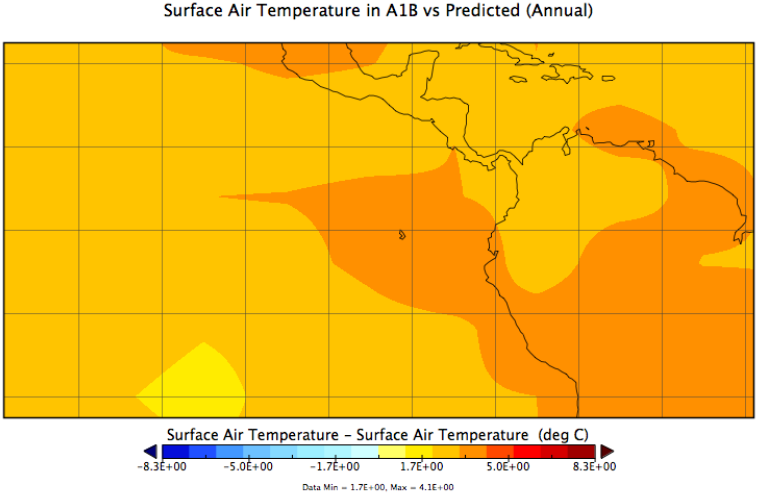
Change in AT (in degrees Celcius) between A1B and Predicted AT above Galápagos annually using a equirectangular map projection.

**Fig. 4.**
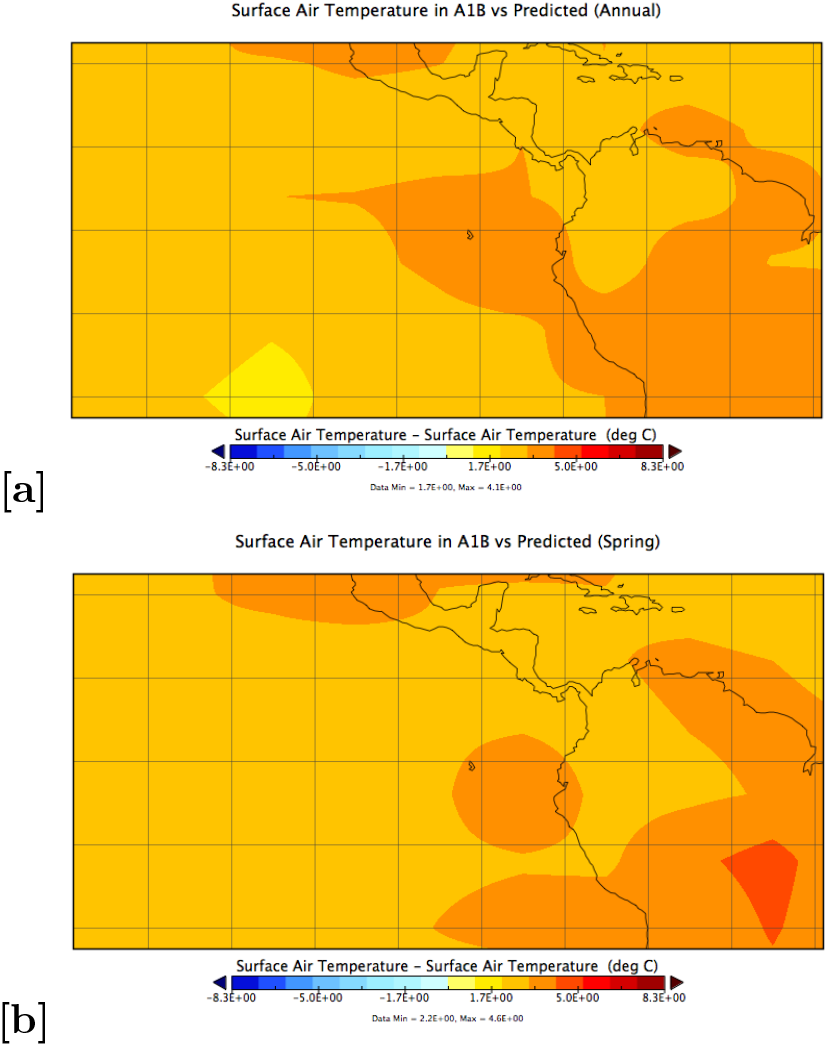
Change in AT (in degrees Celcius) between A1B and Predicted AT above Galápagos in **[a]** winter and **[b]** spring using a equirectangular map projection.

### 3.3 Sea Surface Temperature

Sea surface temperature (SST) is equivalently simple to analyse.

**Fig. 6.**
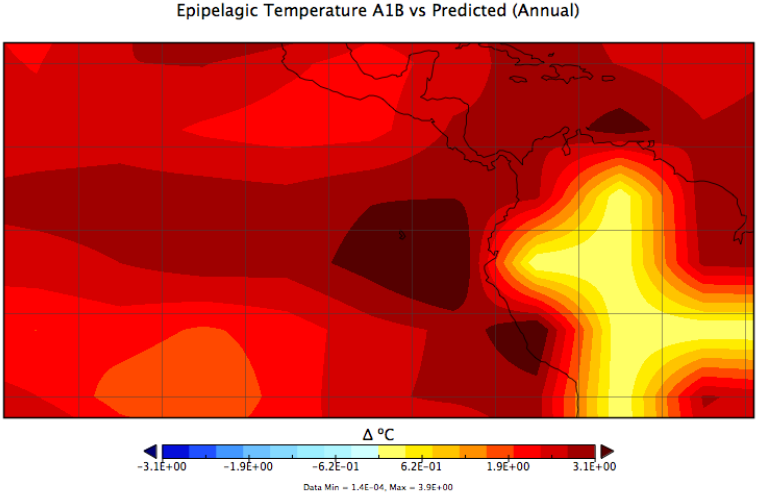
Change in SST (in degrees Celcius) between A1B and Predicted SST around Galápagos annually using a equirectangular map projection.

There is a > 3.1° increase in SST in the A1B scenario. This is true in the annual, autumnal, and hiemal maps.

**Fig. 4.**
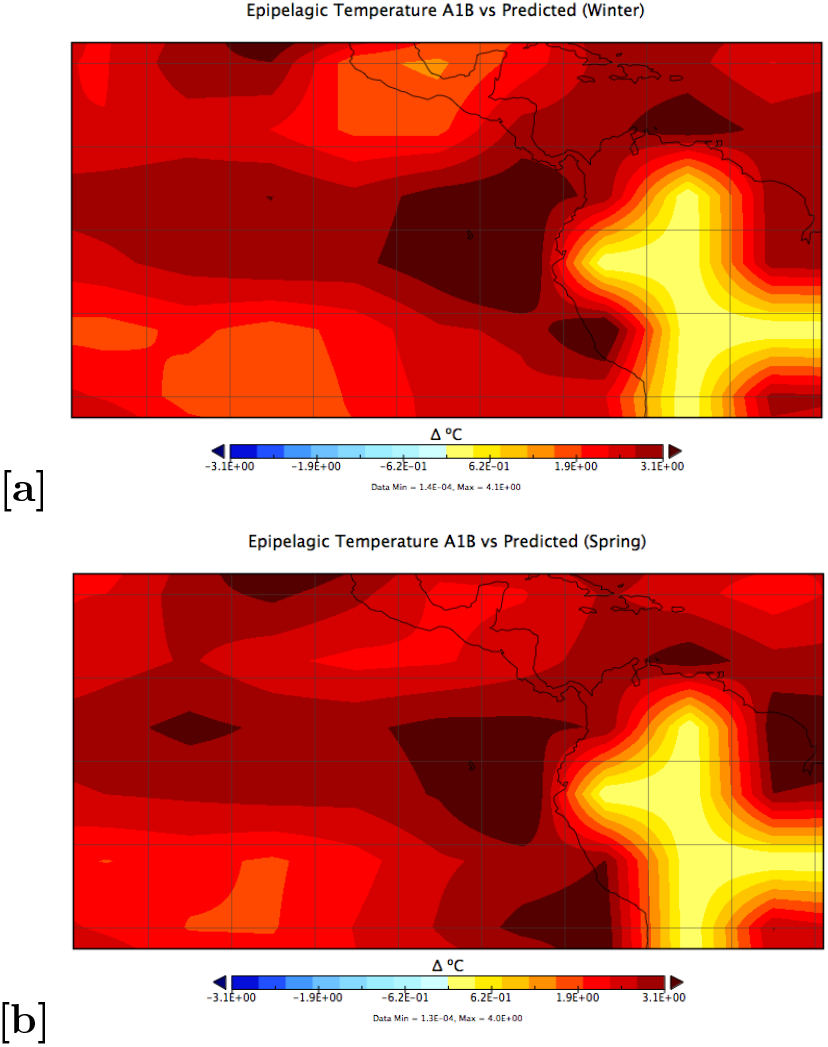
Change in SST (in degrees Celcius) between A1B and Predicted SST around Galápagos in **[a]** winter and **[b]** spring using a equirectangular map projection.

## 4 Discussion

There are many factors that work into figuring out whether *Chelonia mydas agassizii* will be a “winner” or “loser” in an A1B scenario. Since this paper is looking into the effects of climate change on neonate mortality primarily, with an interest in oviposition, fitness, and emergence timings as factors, that is how we will organise our case-by-case analysis.

### 4.1 Ovipositional Effects on Mortality

*Chelonia mydas* oviposition, that is, egg-laying, was originally hypothesised to occur earlier in warmer waters, as it does for sympatrically nesting loggerhead turtles (*Caretta caretta*). However, as Pike (2009) showed, this hypothesis does not stand up to interrogation: *C. mydas* oviposition does not occur earlier in higher SST conditions. However, Hays *et al*. (2002) showed that internesting intervals decrease as SST increases. They plot a regression that shows that at 22°C, internesting intervals approximate 21 days, but as temperatures rise to 28°C, the internesting waiting periods drop to 10 − 14 days.

In the winter of the A1B scenario, 28C SST will measure approximately 28°C, and in the spring, SST will be closer to 30°C (see *Fig. 5a, b*. These correspond to the lower end of the spectrum—10 days or fewer between night crawls. We assume that there must be a lower bound on internesting intervals for females to rejuvenate themselves before repeating their arduous task, and it is unclear what this is at present.

**Fig. 5.**
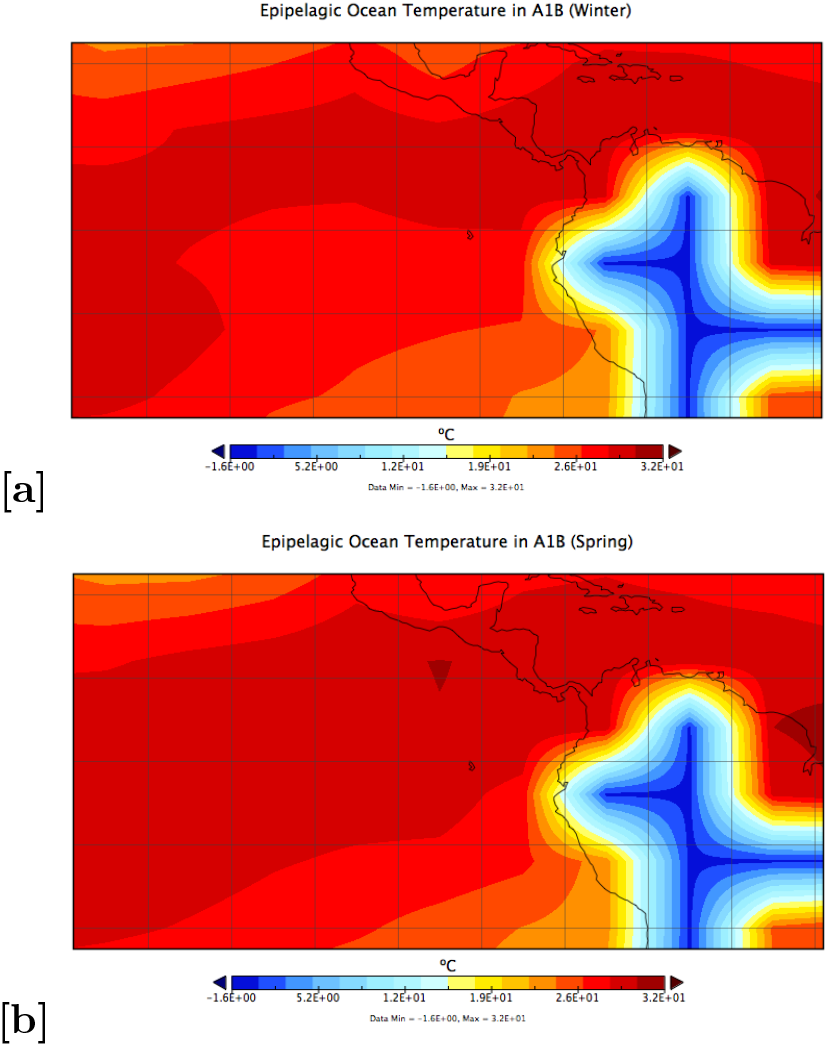
Sea surface temperature (in degrees Celcius) around the Galápagos Archipelago in **[a]** winter and **[b]** spring using a equirectangular map projection.

The effects of this dense period of oviposition could be monumentally helpful in reducing neonate mortality. “There is safety in numbers,” a principle to which sea turtles adhere strictly, and compacted laying season would make this effect even stronger. This could help reduce hatchling mortality significantly. However, if turtles lay at different times during the season, and this effect is well-mixed and random, this effect would be neutralised, meaning that a dense laying season would neither help nor hinder neonates.

### 4.2 Frigate and Reef Fish Predation

One surprising effect of climate change might be increased predation from frigatebirds and reef fish on hatchings. Daytime emergence, as well as emergence during the full, quarter, or three-quarter moons increases reef fish predtion on hatchlings. In fact, the majority of first-year predation occurs in the first hour after entering the water in Australian reef rookeries (Gyuris 1994). Booth & Evans (2011) prove that in reef-surrounded rookeries, reef fish predation is the single largest obstacle to recruiting hatchings. It does not help that green turtle hatchlings have no predator avoidance-system, and must rely on their countershading camoflauge and the power of the frenzy period to survive.

Similarly, Pritchard (IUCN) makes the connection between daytime emergence and frigatebirds predation. According to this article, frigate birds will devour almost every single individual that makes it out onto the sand. My personal experience working at Pacuare, off the East coast of Costa Rica (near Limón) with *Caretta caretta* is that frigatebirds will fight each other over the tiniest morsel, appearing out of nowhere to snatch up your subject.

Pritchard pushes further the connection, however, and states that increased cloud cover results in increased daytime emergence. As shown in *Fig. 6*, there is an fold-change from Predicted SST to A1B of 4%, suggesting that there will be a ≈ 4% increase in frigatebird-related deaths^1^.

**Fig. 6.**
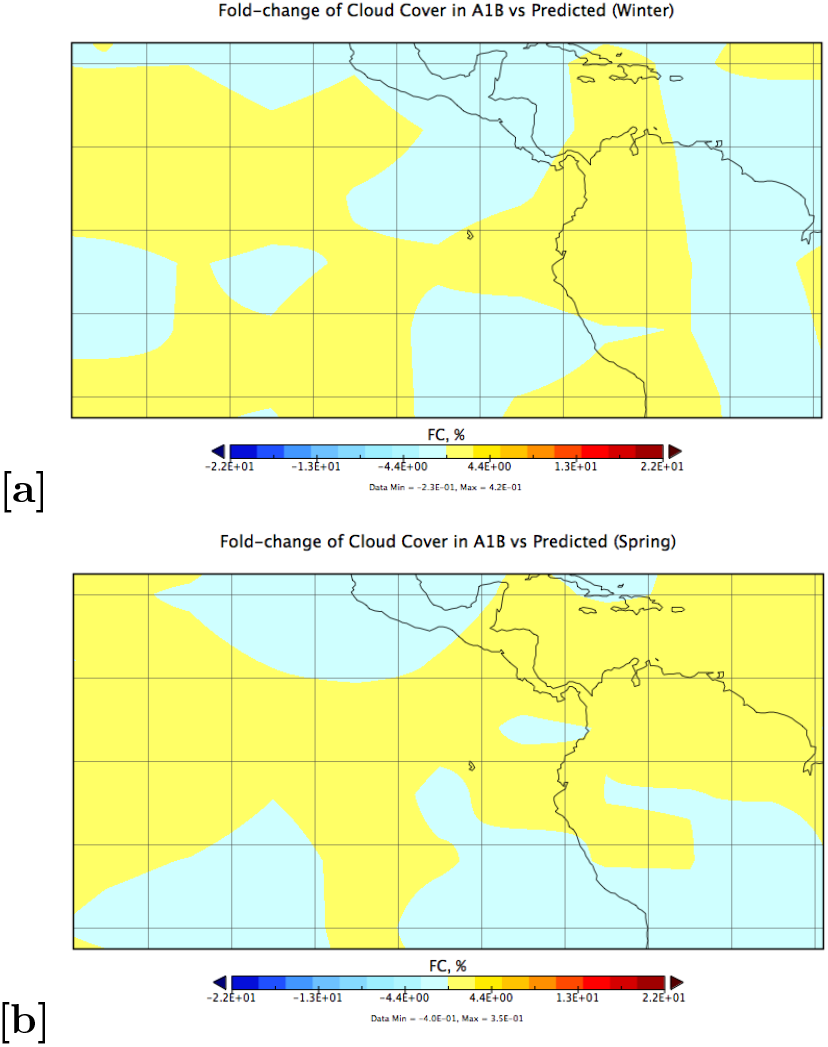
Fold-change (in percentage) of cloud cover above the Galápagos Archipelago in **[a]** winter and **[b]** spring using a equirectangular map projection. This shows a 4% increase in both cases.

### 4.3 Hatchling Fitness and Speed

There has been much discussion on the connection between warm nests and warm waters on speed and fitness of neonates. As discussed in **Frigate and Reef Fish Predation**, reef fish form a significant barrier against a higher percentage of the population from making it past the first hour of their lives. (Whelan and Wyneken (2007) estimate that 5% of all *Caretta caretta* neonate mortality occurs within the first 15 minutes of their active lives.)

There are conflicting views on what connection there is—if any—between nest temperature and hatchling fitness. Ischer *et al*. (2009) claim that colder nests help hatchlings crawl faster, however thrust whilst swimming stays unchanged. They also posit that warmer nests might yield smaller individuals, however it is yet unclear how this might affect their speed^2^. Booth & Evans (2011), however, declare that warmer nests produce slower individuals in general. This paper also puts in the theory that warmer waters help bolster hatchlings to swim faster, helping them escape from the dangers of the reef with greater probability of staying unscathed.

In either case, we may assume that warmer nests have a probability of negatively impacting sea turtle hatchling fitness, and that warmer waters can stimulate neonates to swim faster.

Calculating sand temperature from a large-scale map as we have is not too difficult: Fuentes (2009) and Glen & Mrosovsky (2004) both use AT as a proxy for analysing sand temperature. Fuentes also uses SST as a covariate, but the coefficient here varies greatly from site to site (–0.578 to 1.1231), so we are ignoring it for the sake of simplicity. Air temperature, as shown in the **Results** section, increases by 3 *−* 4°C. This is potentially not ideal for turtle fitness. However, SST increases very significantly, meaning that neonates gain a small speed advantage in A1B. This advantage is negligible in comparison to other impacts discussed in this paper.

### 4.4 Gender Ratio and Thermal Mortality

Nest temperature (as measured through AT as a proxy for sand temperature) has an extraordinary impact on the gender distribution of the occupants of the nest. If the temperature is warmer, 100% of the nest will be female (Glen & Mrosovsky 2004). If the temperature rises still, then the eggs “may experience temperatures that exceed thermal mortality limit” (Fuentes 2009).

Egg-laying females are the limiting factor in population growth for sea turtles, so initially having more females in the population might appear to be advantageous. However, this trend will soon destroy the species, as females will become surplus, and males will disappear entirely. This spells doom for any species, and *C. m. agassizii* is no exception. Additionally, the probability that nests may experience temperatures that exceed the thermal mortality limit is high enough to negate any augmentation of the population the additional females may provide.

Finally, assuming that humans still rely on plastic and fail to properly process it (i.e. create plastic pollution in the ocean), the situation may get worse exponentially faster than previously expected. Washed-up pollution can make a blanket that acts as a greenhouse, heating up the sand even further.

## 5 Conclusion

Under climate change, *Chelonia mydas agassizii* will potentiallty exhibit a denser laying season, which may help reduce neonate mortality. However, frigatebird and reef fish predation will increase in accordance with an uptick in daytime emergence. Hatchling fitness and size or may not decrease as sand temperatures increase, however black turtle hatchlings may swim faster through the warming ocean, potentially bolstering population numbers. Finally, warming nests will produce a surplus of female hatchlings, or an absence of them as the thermal mortality limit is exceeded by global warming and a blanket made of washed-up plastic pollution. It does not appear that Galápagos green sea turtles will “win” in the IPCC’s A1B scenario.

## Footnotes

1 It should be a little smaller than this, given that there may be frigate deaths during the night, however these pale in comparison to the devastating damage done to the population during a daytime emergence fiasco.

2 I am in the process of writing a paper on this topic now with data collected from my frigate-filled adventures at Pacuare.

